# Structural basis for substrate recruitment by AMBRA1 E3 ligase receptor

**DOI:** 10.1101/2022.12.04.519012

**Authors:** Ming Liu, Yang Wang, Fei Teng, Xinyi Mai, Xi Wang, Ming-Yuan Su, Goran Stjepanovic

**Affiliations:** Kobilka Institute of Innovative Drug Discovery, School of Medicine, The Chinese University of Hong Kong, Shenzhen, Shenzhen 518172, China; Department of Biochemistry, School of Medicine, Southern University of Science and Technology, Shenzhen 518055, China; Key University Laboratory of Metabolism and Health of Guangdong, Southern University of Science and Technology,Shenzhen 518055, China

## Abstract

AMBRA1 is a tumor suppressor protein that functions as a substrate receptor of the ubiquitin conjugation system as part of autophagy and cell-cycle regulatory network. The highly intrinsic disorder of AMBRA1 has so far precluded its structural determination. To solve this problem, we analyzed the domain organization and dynamics of AMBRA1 using hydrogen deuterium exchange mass spectrometry (HDX-MS). High deuterium uptake indicates that AMBRA1 is a dynamic and largely unstructured protein, and can be stabilized upon interaction with DDB1, the adaptor of the Cullin4A/B E3 ligase complex. Here we present the cryo-EM structure of AMBRA1 in complex with DDB1 at 3 Å resolution. The structure shows that parts of N- and C-terminal structural regions in AMBRA1 fold together into the highly dynamic WD40 domain, and reveals how DDB1 engages with AMBRA1 to create a binding scaffold for substrate recruitment. AMBRA1 uses its N-terminal helix-loop-helix and WD40 domain to bind the double-propeller fold of DDB1, whereas different regions target the specific cellular substrates for ubiquitination. We also demonstrate that DDB1 binding-defective AMBRA1 mutants prevent ubiquitination of the substrate Cyclin D1 *in vitro* and decreased number of autophagosomes in the cells. Together, these results provide structural insights into AMBRA1-ubiquitin ligase complex and suggests a mechanism by which the AMBRA1 acts as a hub involved in various physiological processes.

## Introduction

The two major intracellular protein degradation pathways in eukaryotic cells, autophagy and the ubiquitin-proteasome system (UPS) are crucial to maintain cell homeostasis and the regulation of several cellular processes, including protein quality control, cell proliferation and apoptosis [1, 2]. Cullin-RING ligases (CRLs) are the largest E3 ligase family in eukaryotes, and are organized by a scaffold protein Cullin and a catalytic RING subunit, RBX1 or RBX2 [1] [4]. Damage-specific DNA binding protein 1 (DDB1) is a component of the Cullin4A/B-RING E3 ubiquitin ligase (CRL4) complex, and functions as an adaptor protein between Cullin4A/B (CUL4A/B), and CUL4A-associated factors (DCAFs) to target substrates for ubiquitination. DDB1 has triple β -propeller (BPA, BPB and BPC) domains and can associate with diverse substrate receptors, which in turn recruits the substrates. Large and mostly unstructured protein AMBRA1 (activating molecule in Beclin-1 regulated autophagy) is one of the DCAF substrate receptors that plays a central role in the communication between autophagy, cell-cycle control and UPS [5]. AMBRA1 can regulate autophagy in several stages. Under starvation conditions, AMBRA1 promotes the ubiquitination of a key autophagy-regulating protein ULK1 by E3 ligase TNF receptor associated factor 6 (TRAF6), results in ULK1 self-association, activation and autophagy induction [6]. In nutrient-rich status, mTORC1 inhibits AMBRA1 by phosphorylation on its Ser-52 [6, 7]. AMBRA1 mediated ubiquitination of ULK1 is prevented through Ser-52 phosphorylation resulting in the termination of the autophagy response. In addition to ULK1, AMBRA1 functions as E3 substrate receptor for Beclin1 ubiquitination during starvation-induced autophagy. In this process, AMBRA1 promotes Lys-63 ubiquitination of Beclin1 by binding to Cullin4 E3 enzyme complex. This modification enhances the interaction of Beclin1 with PI3KC3, which leads to autophagy induction [8]. Recently, AMBRA1 was identified as the major regulator of the stability of D-type cyclins. AMBRA1 limits CDK4/6 activity by mediating ubiquitination and degradation of D-type cyclins as part of the CRL4^DDB1^ E3 ligase complex. Loss of AMBRA1 leads to accumulation of Cyclin D1 and decreased sensitivity to CDK4/6 inhibitors results in increased tumorigenic potential [9-11]. Despite of centrality of AMBRA1 in the intersection between autophagy, UPS and cell growth, the structure and regulatory mechanisms are largely unknown. Here we determined the cryo-EM structure of human DDB1-AMBRA1^WD40^ complex that revealed an unexpected split WD40 domain formed by the N-and C-terminal AMBRA1 regions. HDX-MS analysis show that the binding of DDB1 stabilizes distinct AMBRA1 regions, allowing for structural determination. The N-terminal helix-loop-helix motif of AMBRA1 is essential for DDB1 binding and the point mutations within this region abolished their interaction, leading to prevent the ubiquitination of Cyclin D1 and also reduce the autophagy in cells. Together, these results provide structural insights into AMBRA1-ubiquitin ligase complex and suggests a mechanism by which the AMBRA1 targets proteins involved in essential biological processes.

## Results

### Hydrogen-deuterium exchange analysis of AMBRA1 dynamics

A detailed picture of the mechanism by which AMBRA1 engage CRL4 is required to provide new insight into AMBRA1 regulation, the relationship between autophagy, the UPS pathway and tumor growth, and for development of novel therapeutics. We therefore set out to dissect the molecular basis of AMBRA1 recruitment to CRL4. AMBRA1 has been characterized by WD40 domain in the N-terminal region while the rest of the protein is predicted as intrinsically disordered regions (IDR) (**Fig. 1a**) [12]. Hydrogen deuterium exchange mass spectrometry (HDX-MS) can be used to identify intrinsically disordered regions in a protein or regions with altered conformations between different states [13]. The accessibility of a backbone amide hydrogen is largely dependent on hydrogen bonding and local secondary structure, as well as solvent accessibility [14, 15]. Stably folded secondary structure elements such as α-helices or β-sheets incorporate deuterium to lower degree than interconnecting loops or unstructured regions. In order to differentiate the folded and IDR regions of AMBRA1, purified full length AMBRA1 was analyzed by HDX-MS (**Fig.1b-c**). Experiments were carried out at six timepoints of amide hydrogen exchange (10, 30, 60, 300, 900, 1800 s). A total of 203 peptides covering 98.8% of the sequence were identified and quantified (**Fig. S1, Table S1)**. The backbone amide groups in the first 204 residues of the protein exhibit relative low deuterium uptake, followed by rapid HDX rates characteristic of intrinsically disordered polypeptide segments **(Fig. 1c, S1)**. Besides of the N-terminal region, significantly reduced deuterium uptake was also detected in the AMBRA1 C-terminal region (residues 853-1044), suggesting both regions adopt a folded conformation. Secondary structure and domain prediction of the N-terminal region indicates presence of continuous β-strands that fold into three and half WD40 repeats with length of approximately 40 residues in a single repeat **(Fig. S1)**. The WD40 domains can serve as hotspots for protein-protein or protein-DNA interactions and exhibit a β-propeller architecture, most often comprising seven repeats [16]. Therefore, the AMBRA1 N-terminal region alone was unlikely to exhibit a β-propeller architecture. Thus, we hypothesized that the N-terminal AMBRA1 only forms half of the WD40 domain (WD40-N) and it requires its C-terminal part (WD40-C) to complete the entire β-propeller, connecting by a long loop in between. The C-terminal region (residues 853-1044) was the most probable candidate for the second half of the “split” WD40 domain due to its reduced deuteration levels and the predicted secondary structures of β-strands **(Fig. 1c, S1, S2)**. HDX-MS revealed multiple peptides spanning the predicted AMBRA1 WD40 domain that exhibited bimodal isotope pattern in mass spectra **(Fig. 1d, S2)**. A bimodal distribution could result from EX1 type HDX kinetics where multiple amide hydrogens exchange in the same conformational fluctuation [13, 17]. EX1 kinetic regime is observed in proteins that undergo cooperative unfolding events that simultaneously expose multiple adjacent amide hydrogens for exchange. Under these conditions the refolding rate is much lower than the intrinsic hydrogen exchange rate, and all the amide hydrogens exchange with deuterium in the unfolded (open) state before refolding occurs, resulting in two distinct mass envelopes [18, 19]. The lower mass envelope represents the closed state, and the higher mass envelope represents the open state. To identify the CRL4 binding region in AMBRA1, we co-transfected and reconstituted AMBRA1-CRL4 complex **(Fig. 1b)**. This purified sample was analyzed by HDX-MS and the data were compared to the deuterated mass spectra for apo state of AMBRA1. Upon CRL4 complex binding, the lower mass population becomes more prominent relative to the high mass population in a number of peptides mapping to WD40 domain of AMBRA1 **(Fig. 1d, S2)**. This behavior is exemplified by peptide spanning residues 2-28 **(Fig. S2)**. In AMBRA1 apo form, the closed state is rapidly converted into a more accessible open state. In contrast, CRL4 complex binding significantly stabilized the closed state and slowed down the interconversion between the two states. These changes were consistent with a major stabilization effect on the split WD40 domain dynamics when bound to CRL4 complex, and support its direct role in mediating interaction with E3 ligase adaptors.

**Fig 1:**
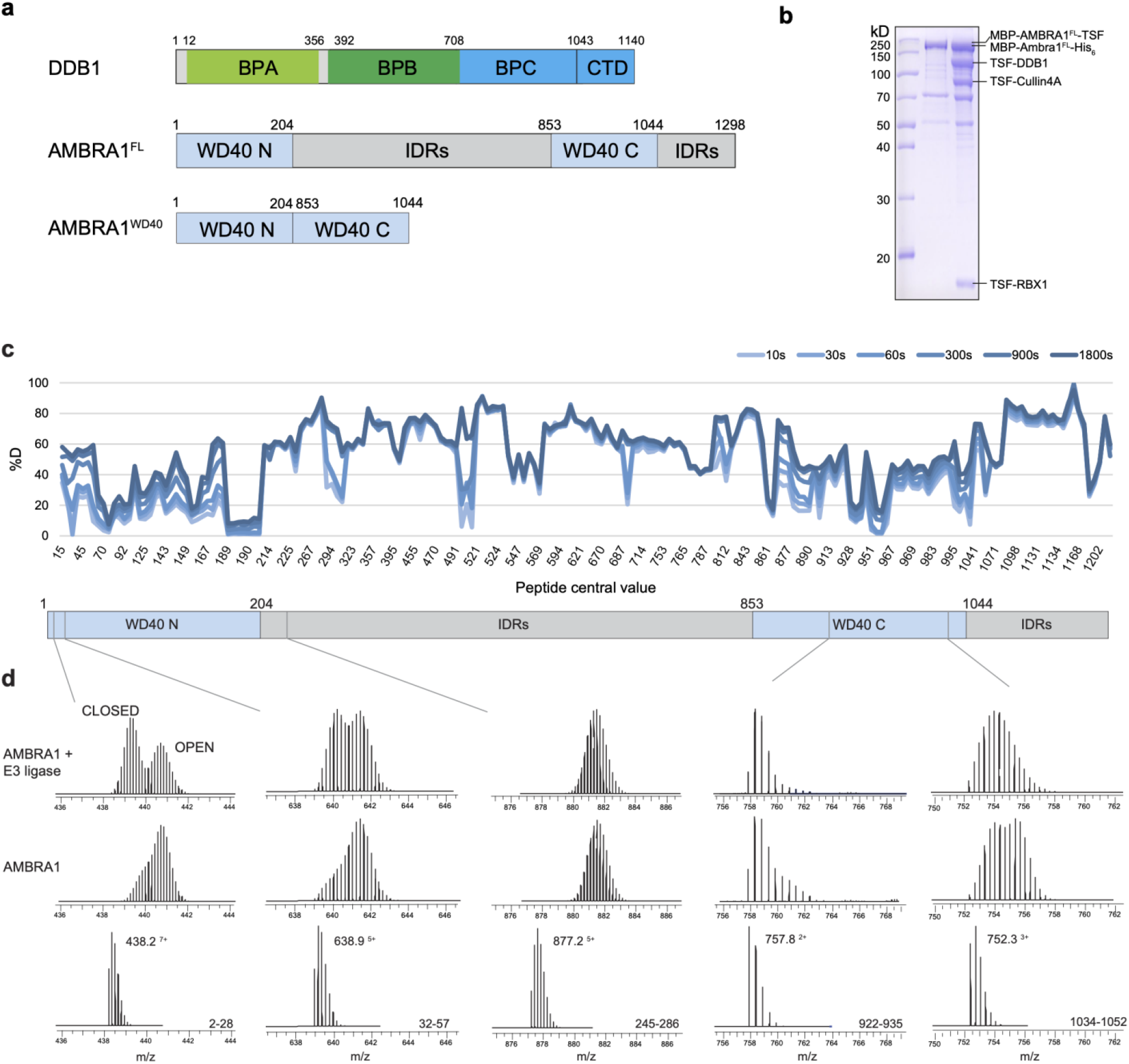
HDX-MS analysis of AMBRA1. a. Annotated DDB1 and AMBRA1 domain schematics. AMBRA1^WD40^ construct for cryo-EM was defined. BPA-BPC, β -propeller A-C; CTD, C-terminal helical domain; FL, full length; IDRs, intrinsically disordered regions. b. The coomassie blue stained SDS-PAGE analysis of purified AMBRA1^FL^ alone and AMBRA1^FL^-DDB1-Cullin4A-RBX1 complex which were used for HDX-MS measurements. TSF, twin-strep-flag. c. Plot representing deuterium uptake by full length AMBRA1, with each data point representing the calculated central residue of an individual peptide. The Y-axis represents % deuteration for a given peptide, at each time point. HDX data statistics is given in Table S1. d. Isotopic envelopes for the selected peptides from AMBRA1, either alone or in E3 ligase complex following 3 s of incubation in D_2_O.

### Cryo-EM structure of the AMBRA1^WD40^ complexed with DDB1

Based on HDX information, a truncated construct comprising residues 1-204 directly fused to residues 853-1044 referred to AMBRA1^WD40^ was designed for cryo-EM structural studies **(Fig. 1a, S3)**, and then co-expressed with DDB1 E3 adaptor in HEK expi293F cells. The resulting complex contained both subunits at apparently equal stoichiometry and the subunits co-migrated as a single peak on gel filtration chromatography **(Fig. 2a-b)**. The complex was enzymatically active as judged by *in vitro* pull-down experiment and ubiquitination assay using Cyclin D1 as a substrate **(Fig.3)**, demonstrating that removal of the AMBRA1 disordered regions between the WD40-N and WD40-C does not disrupt the overall structure and function of the protein.

**Fig 2:**
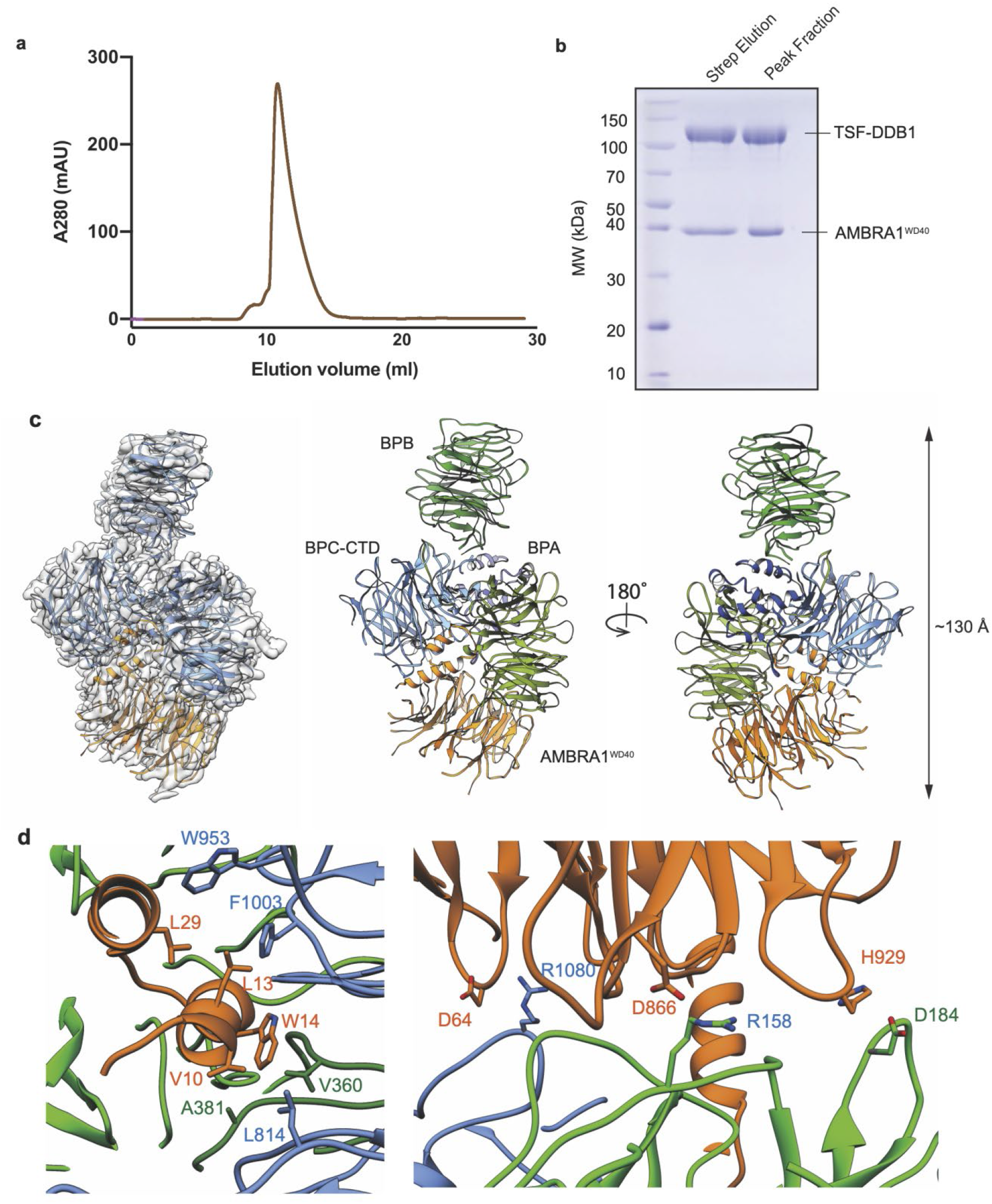
The structural organization of the AMBRA1^WD40^ –DDB1 complex. a. The size exclusion profile (Superdex 200 Increase 10/300) of the purified AMBRA1^WD40^-DDB1 complex. b. The coomassie blue stained SDS-PAGE analysis of AMBRA1^WD40^-DDB1 complex which were used for structure determination. The left lane was the sample from strep elution and the right lane sample was the pooled peak fraction from size exclusion column. c. The cryo-EM density map and the refined coordinates of the AMBRA1^WD40^-DDB1 complex. AMBRA1^WD40^ and the domain of DDB1 were colored as follows: AMBRA1^WD40^, orange; BPA domain, light green; BPB domain, green; BPC-CTD domain, cornflower blue. d. Close-up view between AMBRA1^WD40^-DDB1 interface. The N terminal helix-loop-helix of the AMBRA1 is mainly responsible for the interaction. The key residues contribute to the interface are labelled, some of those residues are further mutated and confirmed with pull down experiment.

**Fig 3:**
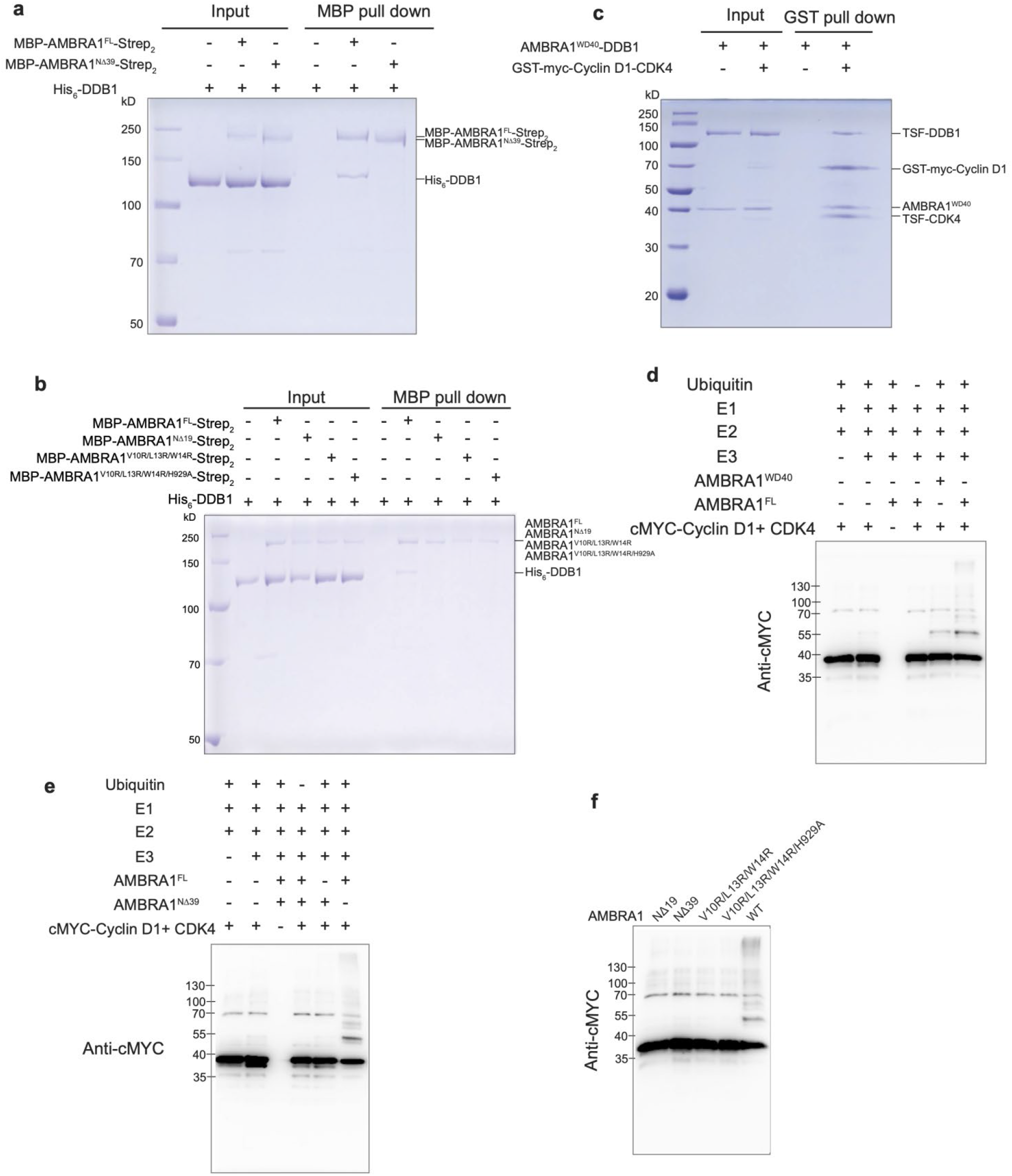
AMBRA1-DDB1-Cullin4A-RBX1 dependent ubiquitination of Cyclin D1. a. *In vitro* pull-down experiment of the full length and N terminal 39 residues deleted AMBRA1 with His^6^ tagged DDB1. The experiment was repeated at least three times and visualized by SDS-PAGE and coomassive blue staining. b. *In vitro* pull-down experiment of the full length, N terminal 19 residues deleted or mutated AMBRA1 with His^6^ tagged DDB1. The experiment was repeated at least three times and visualized by SDS-PAGE and coomassive blue staining. c. *In vitro* pull down experiment of AMBRA1^WD40^-DDB1 with Cyclin D1-CDK4 complex. *In vitro* ubiquitination assay of AMBRA1^FL^/^WD40^ for Cyclin D1. e &f. Ubiquitination assay demonstrated that the N terminal helix-loop-helix of AMBRA1 is responsible for the polyubiquitination of the Cyclin D1. The Ubiquitination assays were analyzed by SDS-PAGE, followed by immunoblot.

We next determined the structure of the AMBRA1^WD40^ complexed with DDB1 by cryo-EM. Cryo-EM images were collected and processed as detailed in the Material and Methods section **(Fig. S3)**. The overall resolution for the structure is around 3.0 Å, allowing to visualize the large side chains in the reconstruction **(Fig. 2c, S4)**. The structure of DDB1-AMBRA1^WD40^ showed 4 individual beta-propeller domains cluster to each other, with the longest dimension of ∼130 Å. DDB1 has three seven-bladed WD40 β-propeller (referred to BPA, BPB and BPC) and a C-terminal helical domain (CTD). BPA and BPC pack against each other in the clamp-shaped double-β-propeller fold, forming a large pocket at the interface. There is a low occupancy in DDB1 as determined by particle imaging analyses and the most flexible part was the BPB domain. 3D classification without alignment analysis demonstrated that this domain can adopt different orientations relative to the BPA-BPC double-β-propeller fold for substrate presentation **(Fig.S5)**. This is consistent with previous studies showing that BPB domain is responsible for interaction with Cullin4 and enables dynamic positioning of the substrates for ubiquitination [20-22].

AMBRA1^WD40^ is characterized by a N-terminal helix-loop-helix motif (E6-K41) and the seven-blade WD40 β-propeller domain which packs against the BPA domain of DDB1 [23]. WD40-N adopts the Blade I-III and the first two strands of Blade IV while WD40-C contributes to the latter two strands of Blade IV and the remaining Blade V-VII. Canonical WD40 domains have seven blades or repeats, each of which contains 40–60 residues, which folds into a β-propeller structure [24]. The WD40 domain of AMBRA1 differs significantly from canonical WD40 domains found in other CRL substrate receptors, such as DCAF1 and CSA. The most prominent difference occurs in the blade IV, at the transition between the βb and βc strand. In DCAF1 the β-hairpin is formed by the βb and βc strands oriented in an antiparallel direction, and linked by a short loop of three amino acids. In AMBRA1, this several-residue long loop is replaced by the 650-residue in length, intrinsically disordered region that separates the WD40-N and WD40-C parts **(Fig. S6a)**.

### AMBRA1 binding to the DDB1 WD40 domain

AMBRA1 interacts with DDB1 via a bipartite structure, consisting of a N-terminal helix-loop-helix motif and a globular core WD40 domain. The helical extension is commonly found in substrate receptors that bind to DDB1 of the CRL4 E3 ubiquitin ligase. The removal of this helical extension (AMBRA1^NΔ39^) completely abolished the interaction with DDB1, demonstrated by *in vitro* MBP pull down assay (**Fig.3a**). The N-terminal helical motif is almost entirely engulfed by the large pocket formed between BPA and BPC double-propeller of DDB1, stabilizing between the hydrophobic interactions contributed by W953, F1003, V360, L814, A381 of BPC domain with V10, L13, W14 in the first helix. Additional DDB1-AMBRA1 interactions include the ionic bonding of AMBRA1 D64 with BPC R1080, AMBRA1 H929 with BPA D184, AMBRA1 D866 with BPA R158 in the loop regions. The AMBRA1 residues are confirmed by mutagenesis by replacement of the hydrophobic residues with positive charged amino acid arginine, further by *in vitro* pull-down assay (**Fig.2d, 3b**). We incubated purified MBP tagged AMBRA1^FL^, AMBRA1^NΔ19^ or AMBRA1^V10R/L13R/W14R^ and AMBRA1^V10R/L13R/W14R/H929A^ with His^6^ tagged DDB1. Only full length AMBRA1 can pull down DDB1, none of the mutants revealed the intermolecular interaction. Collectively, these results confirmed that AMBRA1 N-terminal helix-loop-helix is the most central region for DDB1 association.

Comparison of our AMBRA1-DDB1 structure to other DDB1 complex, the binding between DDB1 and AMBRA1^WD40^ is similar to other adaptors bound to DDB1 including DCAF1(PDB: 6ZUE), Simian virus 5V (PDB: 2B5L), CSA (PDB: 6FCV), DDB2 (PDB: 3EI4), suggesting a common molecular mechanism of DDB1 recognition (**Fig. S6b**). To investigate how AMBRA1 would function in substrates recruitment to the CRL4 E3 ligase machinery, we generated a model of the entire CRL4-AMBRA1 assembly by superimposing the BPB of DDB1 in our structure onto the same domain in CRL4 E3 structure (PDB 2HYE, **Fig.S6c**). Our model revealed a ∼70 Å distance between RBX1 and central cavity of AMBRA1. Notably, the 3D classification without alignment analysis confirmed the plasticity of BPB domain of DDB1, which can adopt diverse orientation to position AMBRA1 for substrate ubiquitination. RBX1 is likely to have reposition as a result of neddylation, therefore establishing an open active conformation to mediating transfer of ubiquitin from E2 to substrate, which shorten the distance between the substrates and the enzymes.

### AMBRA1^WD40^ can directly interact with its substrate Cyclin D1 and mediate its ubiquitination

AMBRA1 can target Cyclin D1 for ubiquitin-mediated degradation, therefore controlling the cell cycle progression through CRL4 E3 ligase [9-11]. To address whether AMBRA1^WD40^ is sufficient for substrate recognition and ubiquitination, we purified Cyclin D1-CDK4 complex for *in vitro* assays. Our GST pull down experiment demonstrated that purified GST tagged Cyclin D1-CDK1 can associate with AMBRA1 ^WD40^ -DDB1 complex (**Fig.3c**). Furthermore, AMBRA1^WD40^ is sufficient to promote ubiquitination of Cyclin D1, but not as robust as full length AMBRA1, indicating the intrinsically disordered region within AMBRA1 may contribute to either augment the interactions or other unknown functions (**Fig.3d**).

AMBRA1 functions as a substrate receptor to recruit Cyclin D1. Therefore, disrupting the interaction between AMBRA1 and DDB1 would fail to mediate the ubiquitination of its substrate. Indeed, Cyclin D1 is polyubiquitinated if every enzyme for the ubiquitination cascade reaction is included. When we excluded either ubiquitin, CRL4 E3 ligase or full length AMBRA1, we did not observe any ubiquitination. In contrast, AMBRA1^NΔ39^ failed to promote the poly-ubiquitination of Cyclin D1 (**Fig. 3e**). We further tested the effect of AMBRA1 mutants including AMBRA1^V10R/L13R/W14R^ and AMBRA1^V10R/L13R/W14R/H929A^ in promoting the ubiquitination of Cyclin D1-CDK4 (**Fig.3f**), none of these mutants execute the activity, indicating the N terminal helix-loop-helix of AMBRA1 is essential for association with CRL4, which in turn mediate the polyubiquitination of Cyclin D1.

### Mutations that disrupt the DDB1-AMBRA1 interaction interface impair autophagy

Given that AMBRA1 also plays pivotal role in regulating autophagy, we further address if disruption of the interaction interface between AMBRA1 and DDB1 influence autophagy *in vivo*. Short interfering RNA (siRNA) oligo ribonucleotide was used to knockdown endogenous AMBRA1 in HEK293 cells as described in Material and methods section. Upon AMBRA1 knockdown, the protein level of LC3-II was significantly decreased by 40% (**Fig.4a**), suggesting downregulation of AMBRA1 resulted in reduction of basal autophagy. The interaction between AMBRA1 and CUL4 E3 ligase has been shown to mediate the K63-ubiquitylation of Beclin1 which results in increased activity of the lipid kinase VPS34 and induces autophagy [8], we then tested the effect of overexpression of wild type and mutated AMBRA1 on autophagy at basal level in HEK293 cells. After AMBRA1 knockdown, we further transfected empty vector, wild type or mutated AMBRA1 to see if they can rescue the function.

**Fig 4:**
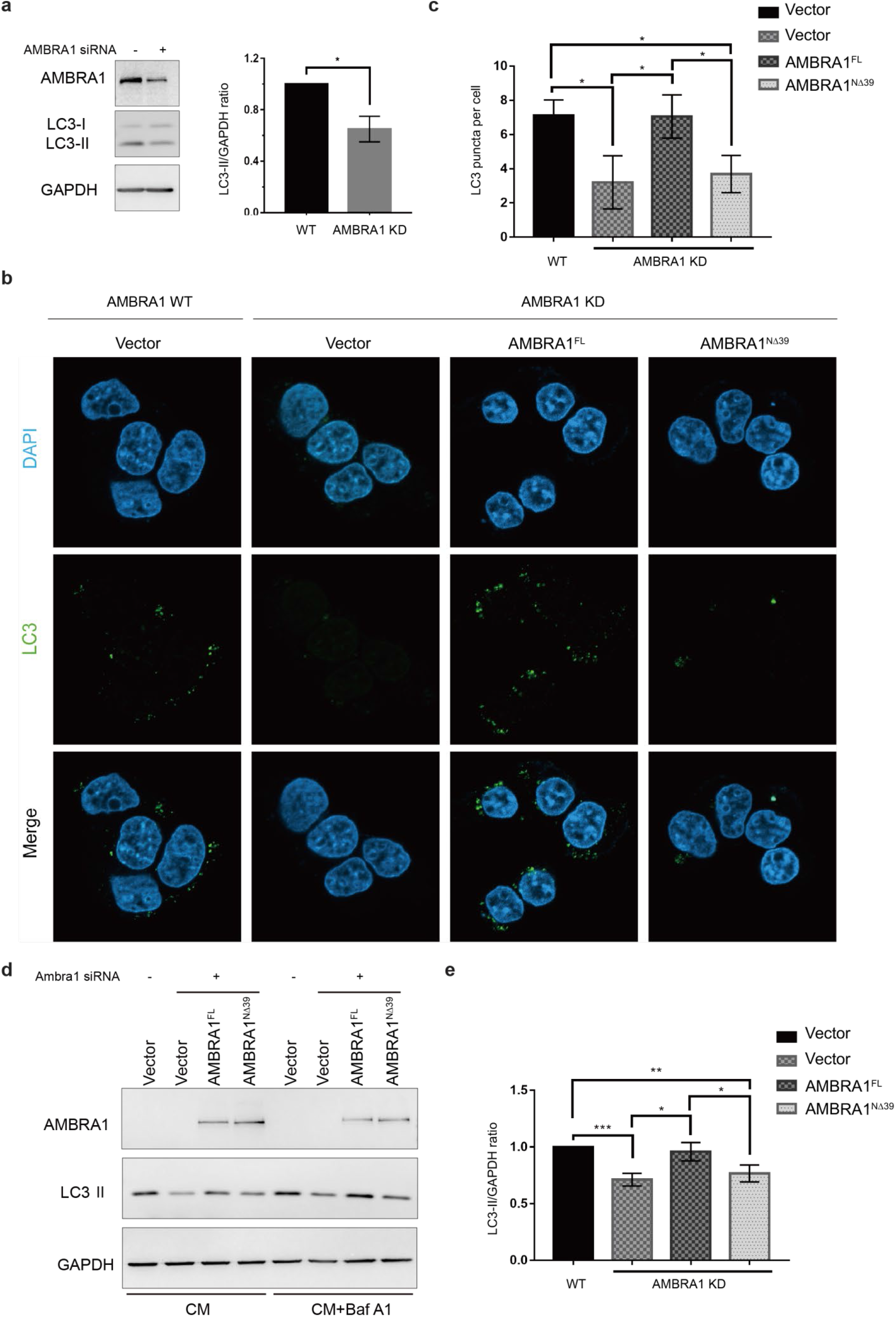
AMBRA1 regulates autophagy. a. AMBRA1 downregulation in HEK293 cells using specific short interfering RNA(siRNA) oligo ribonucleotide. AMBRA1 and LC3 protein levels were analyzed by western blotting (left), GAPDH was used as the protein loading control. Band densitometry was measured by ImageJ, LC3-II levels were normalized to loading control. The quantification of LC3-II/GAPDH ratio from three independent experiments was showed on the right. b. LC3 immunofluorescence assay confirmed that AMBRA1 is essential for the autophagy. After AMBRA1 downregulation, HEK293 cells were transfected with empty vector or WT AMBRA1 or truncated AMBRA1 using X-tremeGENE HP DNA Transfection Reagent (Roche). Cells were treated with 100 nM of bafilomycin A1 for 2 hours at 37 °C before collection. Cells were fixed and stained for LC3 (green) and nuclei (blue) 48 hours after transfection and imaged by confocal microscopy. Representative cells were shown. c. LC3 puncta per cell quantification. LC3 punctum were counted in a total of 50 cells in 8 different fields for each condition. Cells being counted were randomly selected and punctum were counted by using ZEN (Blue edition) software in a blinded fashion by 3 investigators. LC3 puncta per cell in each condition was analyzed using GraphPad Prism 7.0.0 software. Differences between datasets were analyzed using two-tailed statistical tests, the value of p < 0.05 was considered statistically significant and the conventions used to depict the results across this article are as follows: *p ≤ 0.05; **p ≤ 0.01; ***p ≤ 0.001 and Error bars represent standard deviations. d. LC3 immunoblotting confirmed that AMBRA1 is essential for the autophagy. After AMBRA1 downregulation, HEK293 cells were transfected with empty vector or WT AMBRA1 or truncated AMBRA1 using X-tremeGENE HP DNA Transfection Reagent (Roche) according to the manufacturer’s instructions. Where indicated, Cells were treated with 100 nM of bafilomycin A1 for 2 hours at 37 °C before collection. 48 hours after transfection, cells were harvested, AMBRA1 and LC3 protein levels were analyzed by western blotting, GAPDH was used as the protein loading control. e. LC3-II/GAPDH ratio quantification. The LC3-II/GAPDH ratio of cells which were treated by bafilomycin A1 was quantified, Band densitometry was measured by ImageJ, LC3-II levels were normalized to loading control. The quantification of LC3-II/GAPDH ratio from three independent experiments was analyzed using GraphPad Prism 7.0.0 software. Differences between datasets were analyzed using two-tailed statistical tests, the value of p < 0.05 was considered statistically significant and the conventions used to depict the results across this article are as follows: *p ≤ 0.05; **p ≤ 0.01; ***p ≤ 0.001 and Error bars represent standard deviations.

Cells were treated with 100 nM of bafilomycin A1 for 2 hours at 37 °C to block the autophagy flux before collection. Quantifying the number of LC3 punctum from the LC3 immunofluorescence assay, we can see AMBRA1 overexpression can restore autophagy induction while AMBRA1^NΔ39^ overexpression could not, due to AMBRA1^NΔ39^ mutant inability to interact with DDB1. These results show that the interaction between AMBRA1 WD40 domain and CRL4 E3 ligase complex is essential for the induction of autophagy (**Fig.4b-c**). This was further supported by LC3 immunoblotting results showing that AMBRA1 can induce autophagy by interacting with CRL4 E3 ligase. (**Fig.4d-e**). Taken together, these findings indicate the importance of AMBRA1 and DDB1 interaction in autophagy regulation.

## Discussion

This study provides insights into structure and dynamics of AMBRA1, a central hub for coordination of autophagy, cell cycle, cell growth, development and apoptosis [5]. The highly disordered nature and a “split” domain organization of AMBRA1 has largely precluded its structural determination. Intrinsically disordered regions may enable AMBRA1 to perform conformational changes to associate with different interactors simultaneously. So far, AMBRA1 has been reported to recruit and scaffold numerous substrates. For example, its residues 533-751 is the binding site for Beclin1 [25], residues 1075-1077 and 1087-1089 is required for dynein light chain 1 (DLC1) interaction [26], while residues S1043-L1052 are required for association with LC3 and induction of mitophagy [27]. Furthermore, the two PXP motifs (residues P275-R281 and P1206-R1212) were shown to interact with the serine/threonine-protein phosphatase 2A (PP2A) and mediate dephosphorylation and degradation of the proto-oncogene c-Myc resulting in inhibition of cell proliferation and tumorigenesis [28]. These reported regions locate at the intrinsically disordered regions linking WD40-N and WD40-C, and C-terminus. Furthermore, the N-terminal and C-terminal regions of AMBRA1 are involved in interaction with BCL-2, ULK1, ELONGIN B and PARKIN, however the precise interaction sites are not known [29]. In order to gain insights how these regions are organized relative to AMBRA1 WD40 domain in CRL4 structure, we superimposed the Alphafold2 model of AMBRA1 which retains all the loops to our CRL4-AMBRA1 structural model (**Fig. 5**). AMBRA1 WD40 domain is a split domain that must reunite to form a functional substrate receptor. WD40 domain assembly and stabilization by DDB1 may function to organize and bring the diverse substrates and CRL4 E3 ligase towards each other and thus assisting in ubiquitin transfer, resulting in the AMBRA1 ability to coordinate several biological processes in response to a variety of inputs. This mechanism is supplemented by rotational flexibility of DDB1 BPB domain relative to the rest of DDB1-AMBRA1 complex. The DDB1 BPB domain is the attachment site for CUL4A and allows for long-range conformational changes bringing substrates and E2 active site in close proximity. The intrinsically disordered regions contribute to high degree of plasticity in AMBRA1 substrate recognition and recruitment. Several key interactions are mediated by AMBRA1 WD40 domain itself. These include the mTORC1 regulatory phosphorylation site at Ser-52 [6], as well as DDB1 and Cyclin D1 interaction sites. AMBRA1 regulates stability of Cyclin D1 which is commonly upregulated in several cancer, leading to hyperproliferation with genomic instability. AMBRA1 plays a key role in tumorigenesis, and progression, either as an oncogene or as a tumor suppressor [30]. This tightly correlates with a dual role of autophagy in cancer by suppressing the growth of tumors or in tumor growth promotion [31]. The potential therapeutic strategy targeting AMBRA1 will therefore depend from the type of malignancy. AMBRA1-mediated autophagy and apoptosis was associated with chemoresistance in different types of human cancer cells [32-35]. Targeting AMBRA1 expression levels or its regulatory interactions could therefore increase the sensitivity of cancer cells to anticancer drugs or inhibit tumorigenesis and tumor progression. Thus, in providing structural information on AMBRA1, our work may ultimately lead to a better understanding of the molecular basis for regulation of autophagy and apoptosis and to the design of drugs targeting AMBRA1 functional interactions.

**Fig. 5:**
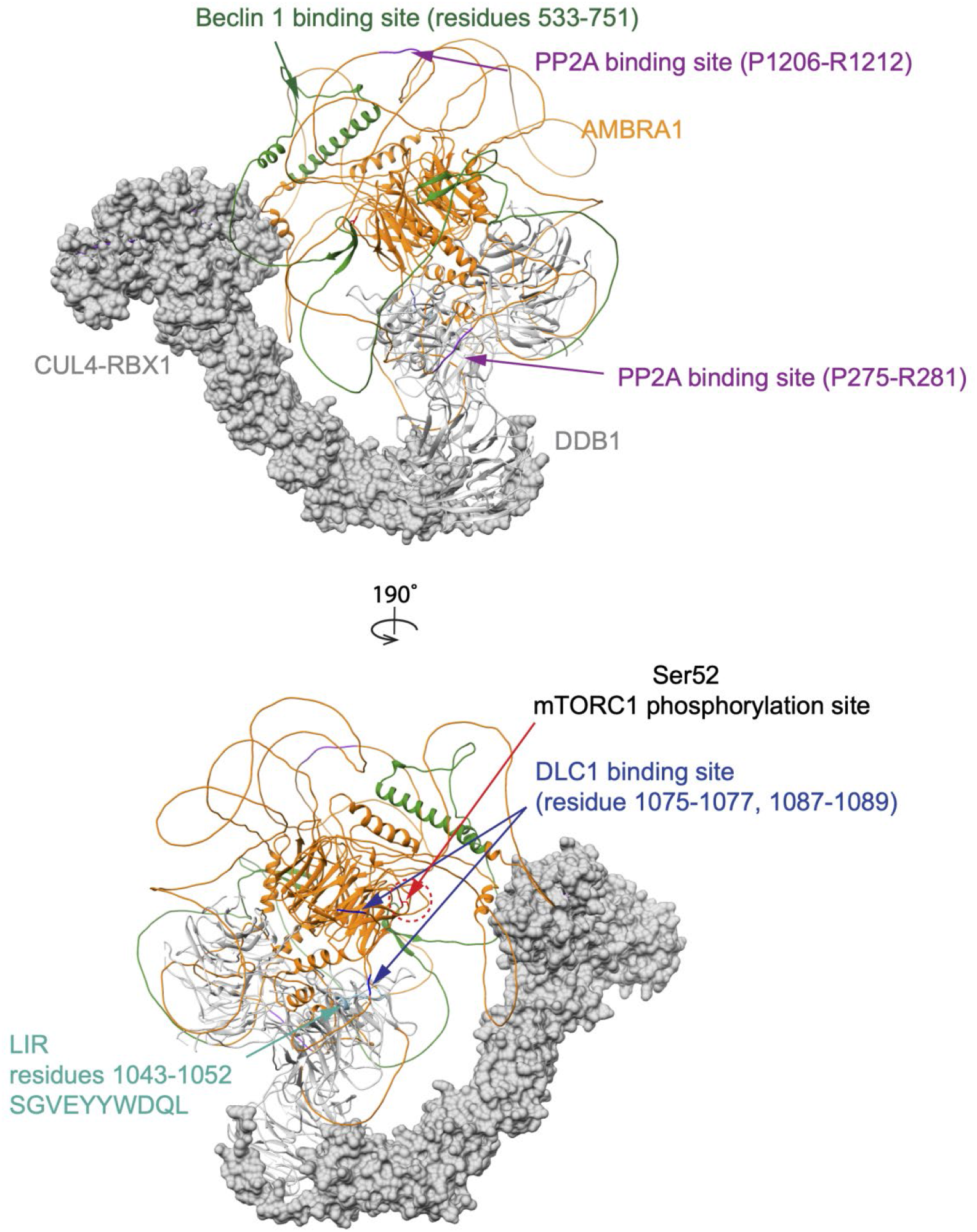
Mapping the interactors binding sites on Alphafold2 model of AMBRA1 which superimposed on Cullin4A-DDB1-RBX1 structure. CUL4 and RBX1 was presented in surface and DDB1 was coloured in gray.

---

**Fig.S1:**
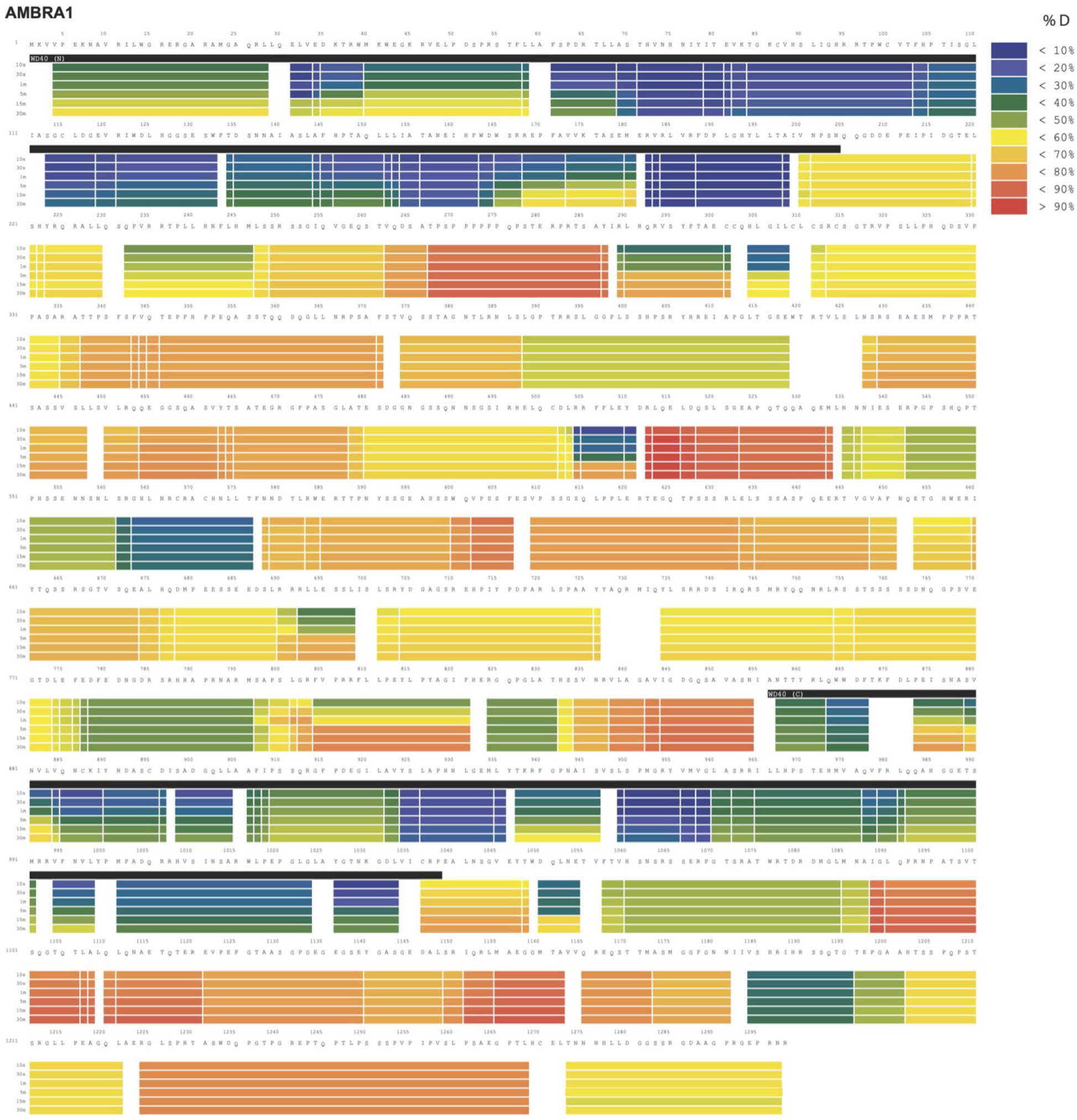
Deuterium uptake data for full length AMBRA1. HDX-MS data are presented in heat map format. Absolute deuterium uptake after 10s, 30s, 1m, 5m, 15 m and 30m are indicated by a colour gradient below the protein sequences. WD40-N and WD40-C regions are indicated above the protein sequence.

**Fig.S2:**
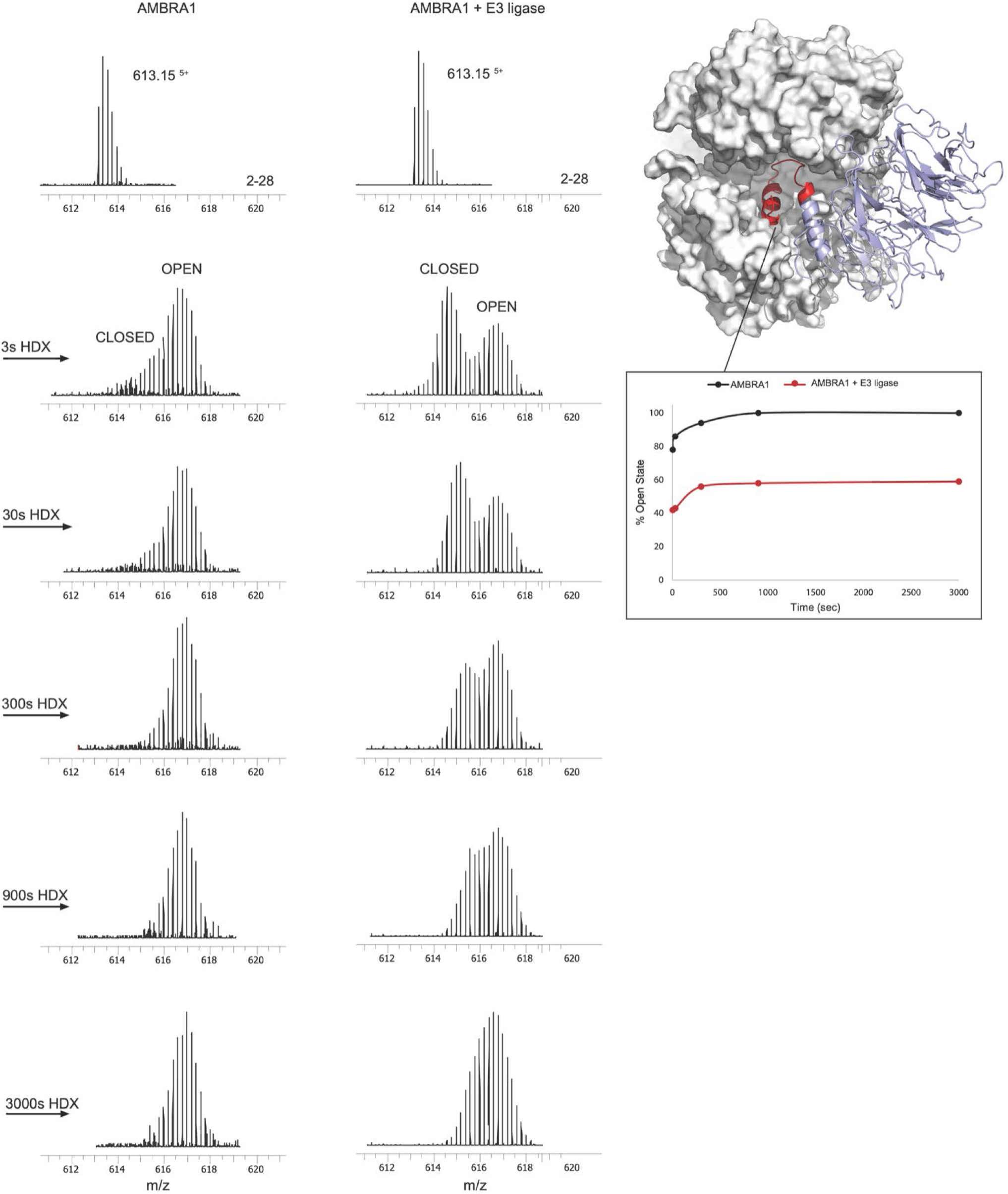
Close to open transition kinetics of AMBRA1 WD40 domain as seen by HDX. Bimodal isotopic envelopes for the AMBRA1 N-terminal helix (peptide spanning residues 2-28), either alone or in E3 ligase complex. Deuteration time points are indicated. Close to open transition kinetics for AMBRA1 helix-loop-helix motif is calculated by fit of two gaussians to the high and low mass subpopulation. Relative amount of the open state is plotted against deuteration time. Structure of the AMBRA1-DDB1 complex interface showing the peptide spanning residues 2-28.

**Fig.S3:**
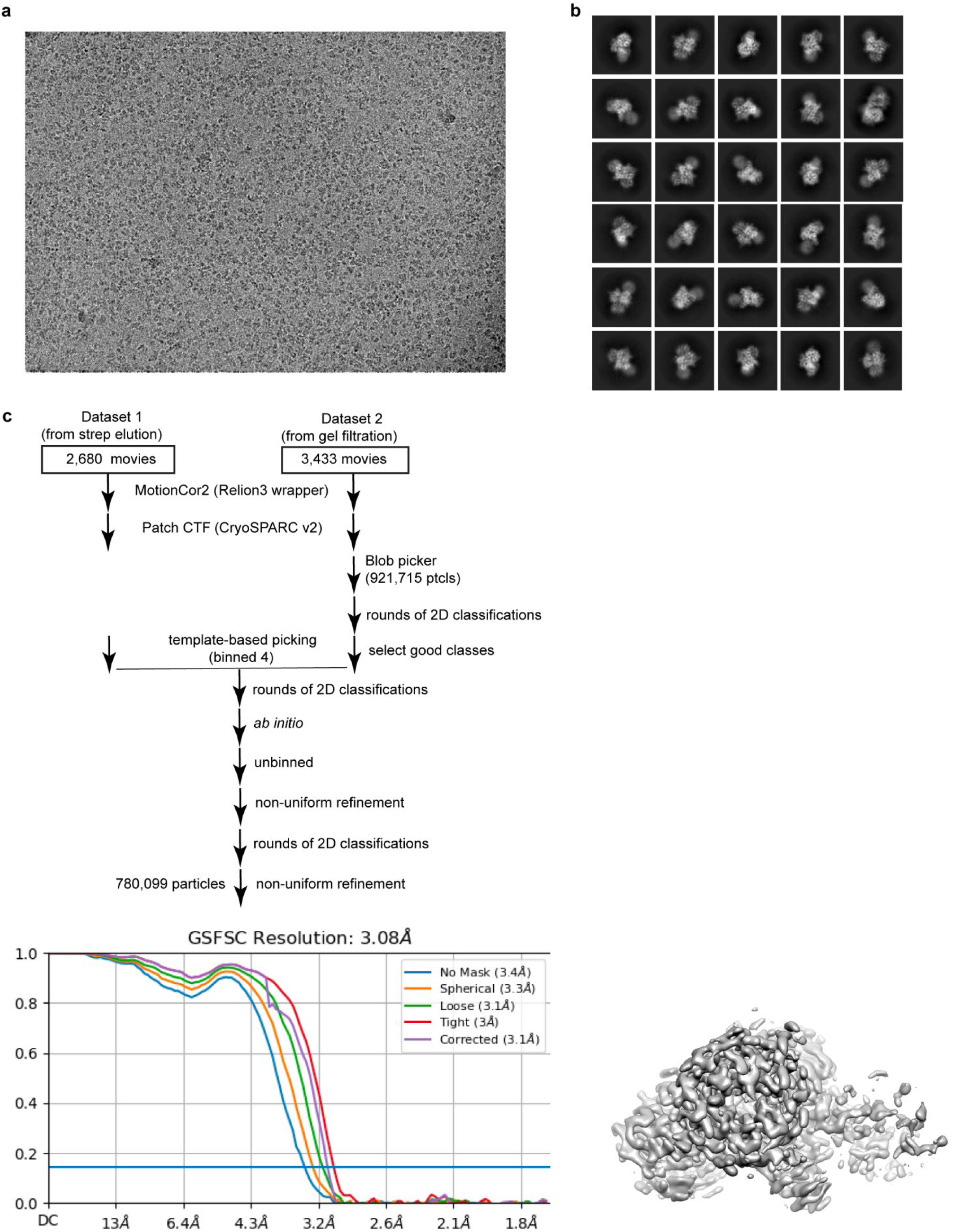
The Cryo-EM data process of the AMBRA1^WD40^ –DDB1 complex. a. A representative motion-corrected cryo-EM micrograph of AMBRA1^WD40^–DDB1 complex. b. Representative 2D class averages for the AMBRA1^WD40^-DDB1 complex. c. Flow-chart of the cryo-EM data processing.

**Fig.S4:**
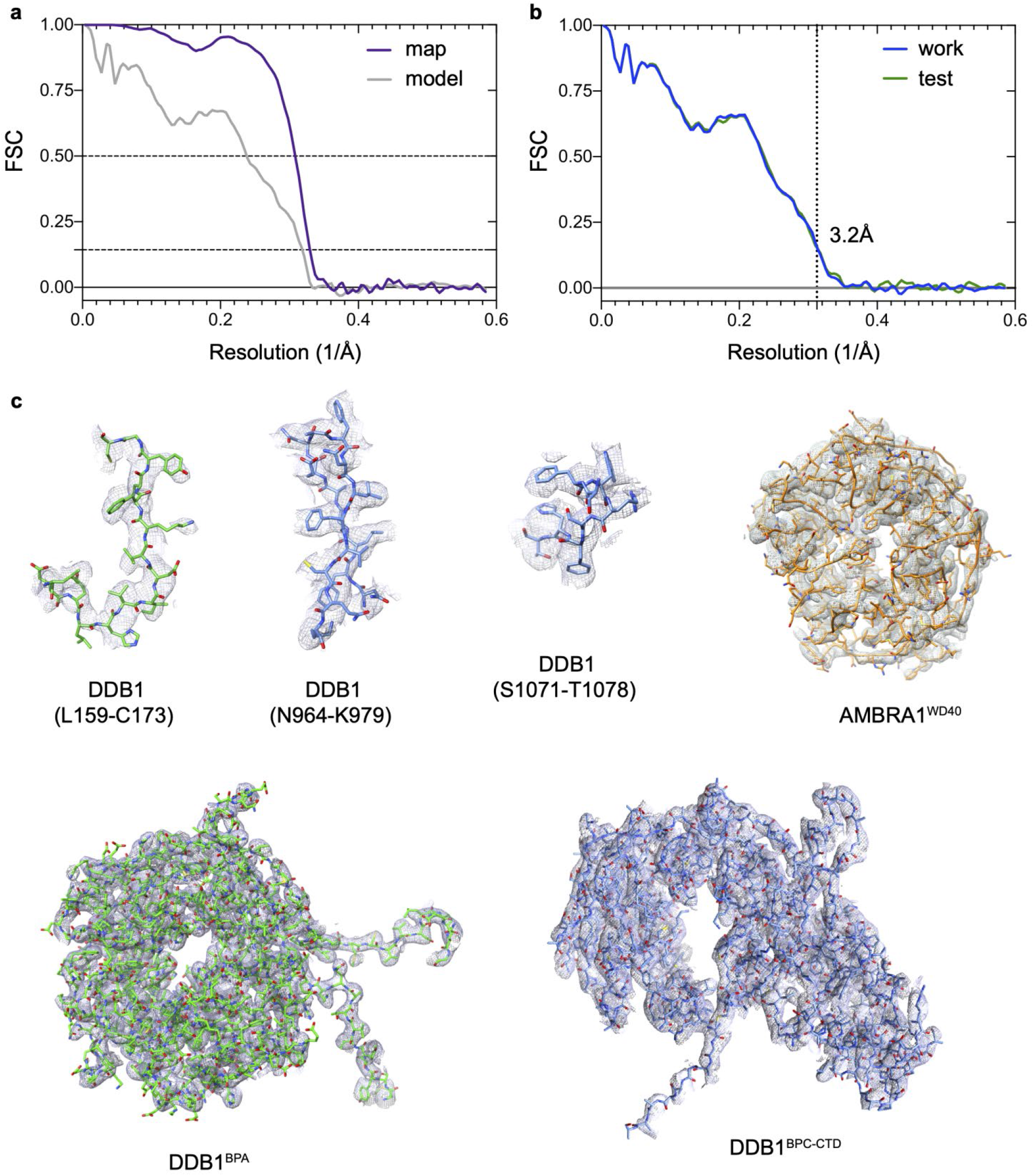
Model building and validation. a, Refinement and map-vs-model FSC. b, Cross-validation test FSC curves to assess over-fitting. The refinement target resolution of 3.2 Å is indicated. c, Refined coordinate model fit of the indicated region in the cryo-EM density.

**Fig.S5:**
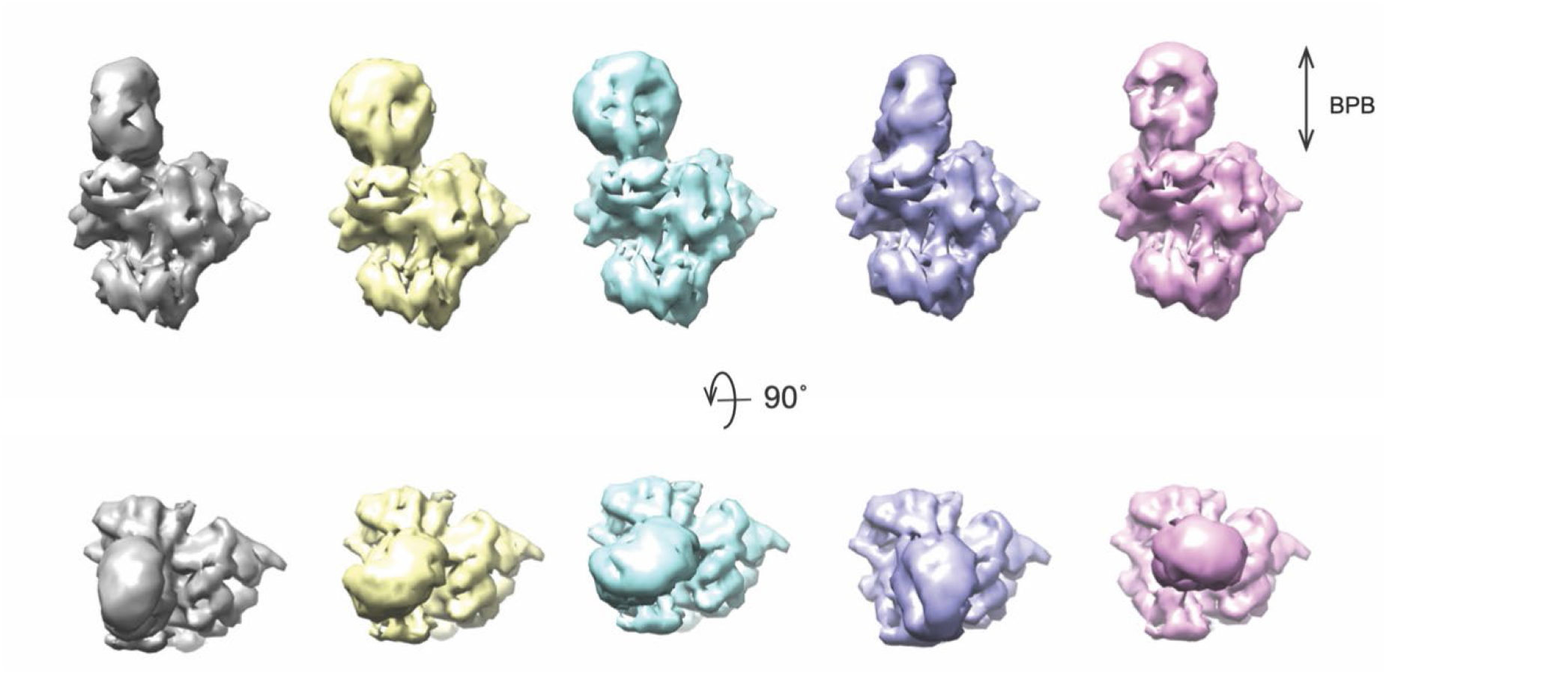
Dynamic of BPB domain of DDB1 that adopts different orientation relative to other domains.

**Fig.S6:**
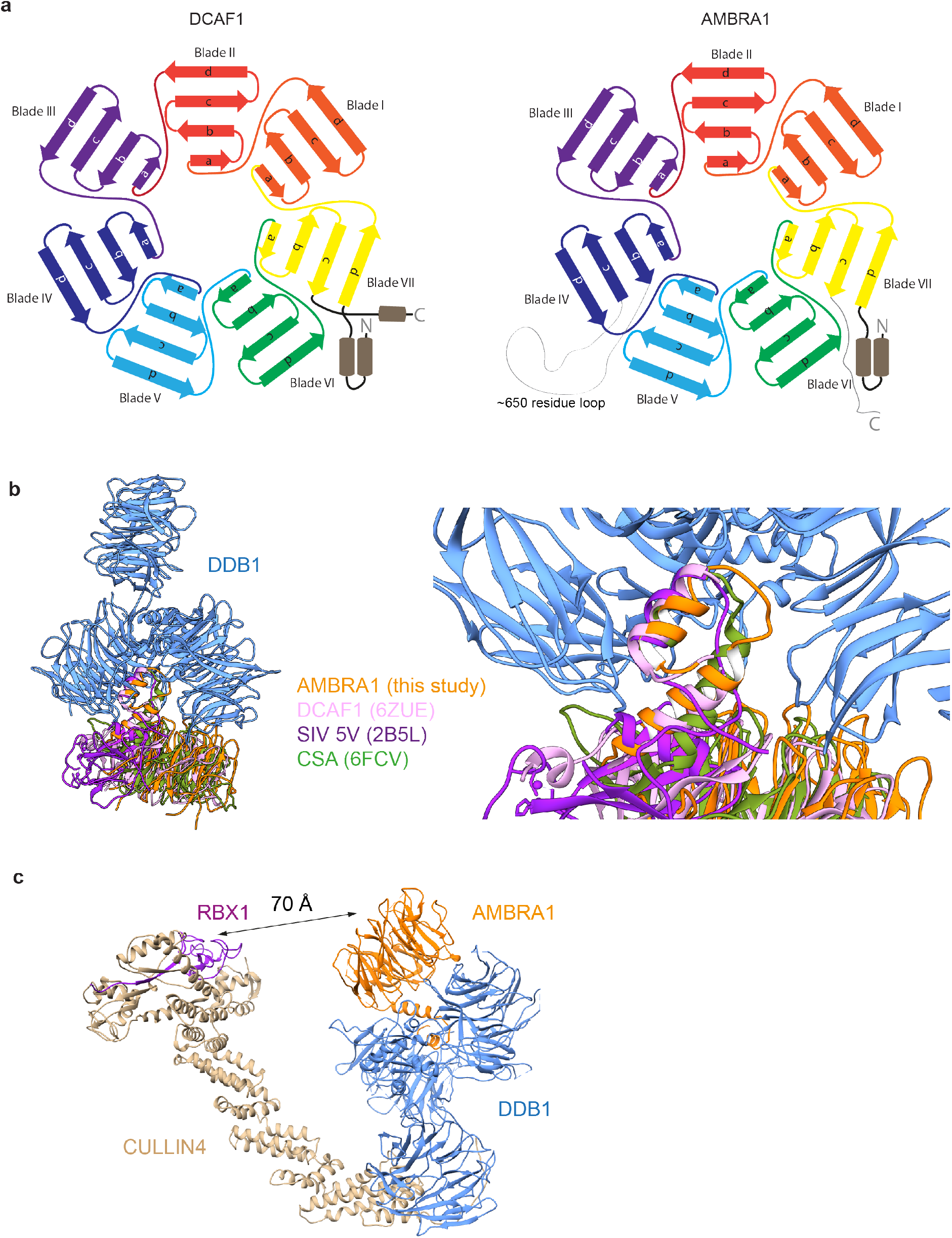
Structural comparison between AMBRA1^WD40^ and other published DDB1 bounded protein. a. Schematic diagram of WD40 domain of DFAC1 (left) and AMBRA1 (right). b. Structural comparion of AMBRA1 (colored in orange, this study), Simian virus 5V (colored in margenta, PDB 2B5L) and CSA (colored in green, PDB 6FCV) bound to DDB1. c. Model of the entire CRL4-AMBRA1 E3 complex.

**Fig.S7:**
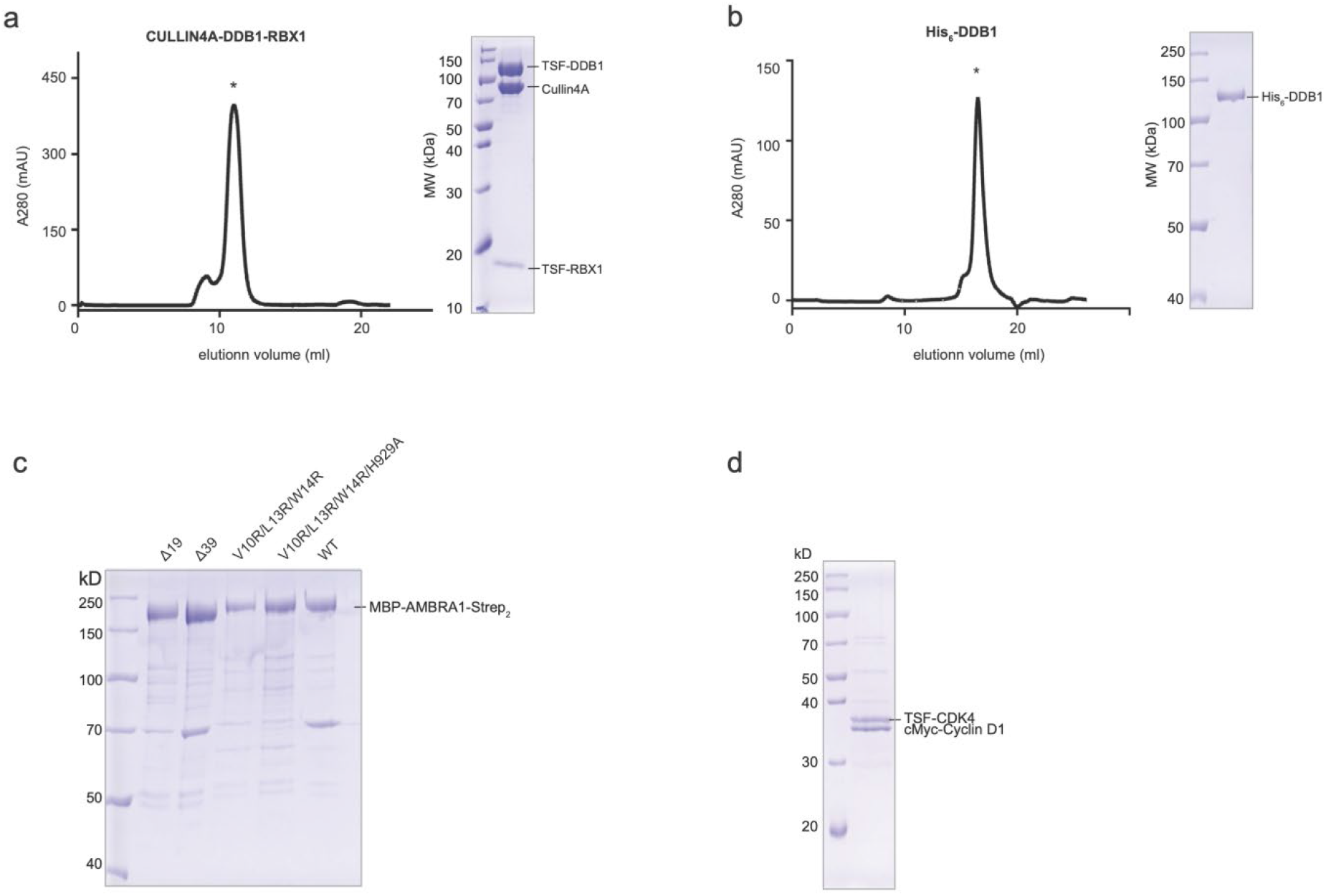
Purification of CRL4 E3 ligase, DDB1, WT/truncated/mutated AMBRA1 and Cyclin D1-CDK4 complex. a. The size exclusion profile (Superdex 200 Increase 10/300) and the coomassie blue stained SDS-PAGE analysis of the purified protein Cullin4A-DDB1-RBX1(E3) ligase which were used for *in vitro* ubiquitination assay. b. The size exclusion profile (Superose 6 Increase 10/300) and the coomassie blue stained SDS-PAGE analysis of the DDB1 which was used for MBP pull down assay. c. The coomassie blue stained SDS-PAGE analysis of the WT/truncated/mutated AMBRA1. d. The coomassie blue stained SDS-PAGE analysis of the Cyclin D1-CDK4 complex.

## Materials and Methods

### Cloning and mutagenesis

The gene for full length AMBRA1 (residue M1-R1298) was codon optimized. The genes encoding DDB1, Cullin4A, RBX1, Cyclin D1 and CDK4 were amplified from cDNA by PCR and subcloned into pCAG vectors with different tags using KpnI and XhoI cutting sites, respectively. For full length AMBRA1 used in in vitro experiments, it was constructed as a N-terminal MBP tag with a His^6^ tag in the C terminus. AMBRA1^WD40^ (residues 1-204/853-1044) was subcloned with N-terminal GST tag followed by a TEV cutting site. For Cyclin D1, it was cloned with N-terminal GST-TEV-myc tag. DDB1, Cullin4A, RBX1 and CDK4 were constructed with N terminal Twin-Strep-FLAG (TSF) tag. The AMBRA1 mutants were generated by PCR-based site-directed mutagenesis approaches. For AMBRA1^FL^ (residue M1-R1298), AMBRA1^NΔ19^ (residue A20-R1298), AMBRA1^NΔ39^ (residue M40-R1298), AMBRA1^V10R/L13R/W14R^ and AMBRA1^V10R/L13R/W14R/H929A^ used in the MBP pull down experiment, they were cloned as a N-terminal MBP tag and a Twin-Strep tag in the C terminus. DDB1 was also subcloned into pFastBac-dual vector with a His^6^ tag in the N terminus for insect cell expression. For in vivo cell experiments, taggless AMBRA1^FL^, AMBRA1^NΔ19^, AMBRA1^NΔ39^, AMBRA1^V10R/L13R/W14R^ and AMBRA1^V10R/L13R/W14R/H929A^ were cloned into pCAG vector.

### Protein expression and purification

For mammalian cell expression, N-terminal GST tag human AMBRA1^WD40^ and N-terminal TSF DDB1 were co-expressed for structural studies, N-terminal GST myc tag human Cyclin D1 and N-terminal TSF tag CDK4 were co-expressed for in vitro ubiquitination experiment. Full length MBP tagged AMBRA1 with C-terminal His^6^ tag was co-expressed with TSF tagged Cullin4A-DDB1-RBX1 E3 ligase for HDX measurement. MBP tagged full length AMBRA1, AMBRA1^NΔ19^, AMBRA1^NΔ39^, AMBRA1^V10R/L13R/W14R^ and AMBRA1^V10R/L13R/W14R/H929A^ with a C-terminal Twin-Strep tag were also expressed individually in Expi293F cells for in vitro pull-down experiment. N-terminal His^6^ tag human DDB1 was expression in sf9 cell.

The Expi293F cells were grown in Union-293 medium and used for protein expression, with 1 mg DNA and 4 mg PEI per 1 liter of cells at density of 2x10^6^ cells/ml. Cells were harvested after 3 days and washed with 1X cold PBS.

For AMBRA1^WD40^ and DDB1 coexpression, the cell pellet was lysed in lysis buffer (50 mM HEPES pH 7.4, 200 mM NaCl, 10 % glycerol(v/v), 1% Triton-X100 (v/v), 2 mM MgCl_2_, 5 mM beta-mercaptoethanol) with proteases inhibitors (1mM PMSF, 0.15uM Aprotinin, 10uM leupeptin, 1uM pepstain) for 20 min at 4 °C. After centrifugation at 18,000 rpm for 30 min, GST beads were incubated with the cell supernatant at 4 °C for 2 h. The beads were washed three times with wash buffer (50 mM HEPES pH 7.4, 200 mM NaCl, 2 mM MgCl_2_, 5 mM beta-mercaptoethanol). Bound proteins were eluted with wash buffer containing 50 mM reduced glutathione. After TEV digestion overnight, the elution was then subjected to Streptactin sepharose resin, washed three times with wash buffer and eluted with gel filtration buffer (50 mM HEPES pH7.4, 200 mM NaCl, 2 mM MgCl_2_ and 1 mM TCEP) supplemented with 5 mM desthiobiotin. The protein elution was further purified by Superdex 200 Increase 10/300 GL (GE Healthcare). Cyclin D1-CDK1 complex were purified in the same procedure as described above.

For full length AMBRA1 and Cullin4A-DDB1-RBX1 E3 ligase coexpression, amylose beads were incubated with the cell supernatant at 4 °C for 1 h. The beads were washed three times with wash buffer and eluted in wash buffer containing 10 mM maltose. The elution was further applied to Streptactin sepharose resin, washed three times and eluted with gel filtration buffer containing 5 mM desthiobiotin. Full length AMBRA1^NΔ19^, AMBRA1^NΔ39^, AMBRA1^V10R/L13R/W14R^ and AMBRA1^V10R/L13R/W14R/H929A^ was purified in the same protocol.

The sf9 insect cell was grown in ESF921 medium and infected with recombinant baculovirus (1 to 50 v/v ratio) at density of 2x10^6^ cells/ml. Cells were harvested after 3 days and washed with 1X PBS. For DDB1 purification, the cell pellet was lysed in lysis buffer (50 mM Tris pH 8.0, 200 mM NaCl, 20 mM imidazole, 1% Triton-X100 (v/v), 2 mM MgCl_2_, 5 mM Beta-mercaptoethanol) with protease inhibitors for 20 min at 4 °C. After centrifugation at 18,000 rpm for 30 min, the supernatant was passed over Ni-NTA affinity beads. The beads were washed three times with wash buffer (50 mM Tris pH8.0, 200 mM NaCl, 20 mM imidazole, 2 mM MgCl_2_, 5 mM beta-mercaptoethanol) and eluted with elution buffer (50 mM Tris pH 8.0, 200 mM NaCl, 200 mM imidazole, 2 mM MgCl_2_, 5 mM beta-mercaptoethanol). The protein elution was further purified by Superose 6 Increase 10/300 GL (GE Healthcare) in gel filtration buffer (50 mM HEPES pH 7.4, 200 mM NaCl, 2 mM MgCl_2_, 1mM TCEP). The protein sample was flash frozen in liquid nitrogen and stored at -80 °C until use. All purification procedures was carried out at 4 °C. The proteins used in this study are shown in Fig. S7.

### Baculovirus generation

For insect cell expression, target gene was cloned into pFastBac-dual vector, then transformed into DH10Bac Competent cell. The Bacmid was extracted and used to transfect the insect cell using Cellfectin II Reagent (Gibco) in 6-well plate, according to the manufacturer’s instructions. 4 hours after transfection, replace medium with fresh ESF921 medium containing 10% fetal bovine serum and 1% Antibiotic-Antimycotic solution. Cells were incubated at 27 °C for 4 to 5 days, then collect the supernatant and centrifuge at 2500rpm for 5 minutes to remove cells and large debris, transfer the supernatant into new stelized tube, that’s P0 virus. To generate P1 virus, prepare 50 ml Sf9 cell with 2 × 10^6^ cells/ml density, add 2ml P0 virus into the cell, wait for 4-5 days and check the viability to drop to ∼50-60%, Harvest the cell by centrifuge at 2500rpm for 15 min, Collect the supernatant, that’s P1 virus. add FBS to a final concentration of 5% and store at 4 °C prevent from light. To generate P2 virus, follow the same procedure, the amount of P1 virus being used depending on the viral titers. For long-term storage, aliquot and store at -80 &C for later use.

### Hydrogen – deuterium exchange mass spectrometry (HDX)

H/D exchange was carried out by 10-fold dilution of 0.5 µM AMBRA1 or AMBRA1-E3 ligase complex into a D_2_O buffer (20 mM HEPES pH 7.5, 150 mM NaCl, 2 mM MgCl) to a final volume of 100 µL. After different time periods of incubation at 25 °C, the reaction was quenched by addition of ice-cold quench solution (1:1, v/v) containing 2M guanidinium hydrochloride and 0.2M citric acid. This reaction was digested on an immobilized pepsin column inside a manual HDX system with the temperature maintained at 0 °C. Eluted peptides were desalted using chilled trap column (1mm × 15mm, Acclaim PepMap300 C18, 5μm, Thermo Fisher Scientific) for 5 min at a flowrate of 200μL/min and 0.1% formic acid as mobile phase. Subsequent peptide separation was performed on the chilled ACQUITY BEH C18 (2.1 × 50mm) analytical column using a first gradient ranging from 9 to 45 % of buffer B (80% acetonitrile and 0.1% formic acid) for 10 min followed by a second gradient ranging from 45 to 99% of buffer B for 1 min, at an overall flow rate of 50μl/min. Peptides were ionized via electrospray ionization and analyzed by Orbitrap Eclipse and QExactive HFX (Thermo Fisher) mass spectrometer. Non-deuterated samples peptide identification was performed via data dependent tandem MS/MS experiments and analyzed by Proteome Discoverer 2.5 (Thermo Fisher). Mass analysis of the peptide centroids was carried by HDExaminer v3.3 (Sierra Analytics, Modesto, CA), followed by manual verification for each peptide. No corrections for back exchange that occurs during digestion and LC separation were applied. For peptides exhibiting a bimodal distribution each time point is fitted with two Gaussian peaks with different means and areas but with similar width into the spectra. Next, the fitted peak area parameter was used to calculate the relative amount of open state by taking the ratio of high and the sum of high and low mass subpopulation.

### Cryo-EM grid preparation and data acquisition

Two samples of human AMBRA1^WD40^-DDB1 complex were prepared for cryo-EM data acquisition. The first one is the protein elution from strep beads while the other one is the peak elution from size exclusion column. For cryo-EM grid, 3 μl of 0.15-0.2 mg/ml protein sample were deposited onto freshly glow-discharged Quantifoil R1.2/1.3 Cu300 mesh grids and plunged into liquid ethane using a FEI Vitrobot Mark IV after blotting for 3 sec with blot force 0 at 4°C and 100% humidity. A total of 6,147 movies were collected on a Titan Krios electron microscope operating at 300 kV equipped with Gantan K3 camera at a defocus of -1 μm to -1.8 μm in counting mode, corresponding to a pixel size of 0.85 Å. Automated image acquisition was performed using SerialEM with a 3x3 image shift pattern. Movies consists of 50 frames, with a total doses of 58.96 e^-^/Å^2^, with a total exposure time of 3 sec and a dose rate of 14.2 e-/pixel/sec. Imaging parameters for the dataset are summarized in Table S2.

### Cryo-EM data processing

The first dataset included 3,456 movies and the another one has 2,691 movies. For each dataset, the movies were firstly aligned using MotionCor2 wrapper in Relion 3 to correct the specimen movement and then imported into cryoSPARC v2. CTF fitting and estimation were performed by patch ctf estimation. After particle picking with blob picker in the second dataset, 921,715 particles were picked and subjected to 2D classification. The good classes were selected as templates for the entire datasets, which generating 2,082,649 particles. The particles were cleaned up with iterative 2D classification and subjected for ab initio and non-uniform refinement, resulted in 3.0 Å resolution map. 780K particles were imported into Relion 3.1 for 3D classification without alignment analysis. The map was postprocessed by deepEMhancer with tight target modes for model building. All reported resolutions are based on the gold-standard FSC 0.143 criterion.

### Atomic model building and refinement

The coordinates for DDB1 (PDB: 2B5M) and AMBRA1^WD40^ generated from Alphafold structure prediction were rigid body fitted separately into the density map using UCSF Chimera. Atomic coordinates were refined by iteratively performing Phenix real-space refinement and manual inspection and correction of the refined coordinates in Coot. To avoid overfitting, the map weight was set to 2 and secondary structure restraints were applied during automated real-space refinement. Model quality was assessed using MolProbity and the map-vs-model FSC. A half-map cross-validation test showed no indication of overfitting. Figures were prepared using UCSF Chimera version 1.15. The cryo-EM density map has been deposited in the Electron Microscopy Data Bank under accession code XXXXX and the coordinates have been deposited in the Protein Data Bank under accession number XXXX.

### MBP pull down assay

0.5 μM of purifed AMBRA1^FL^, AMBRA1^NΔ19^, AMBRA1^NΔ39^, AMBRA1^V10R/L13R/W14R^ and AMBRA1^V10R/L13R/W14R/H929A^ was incubated with 2.5 μM of purified DDB1 in 500 uL reaction volume supplemented with 50 μL amylose beads respectively. After 2 hr incubation at 4 °C, the beads were washed three times with gel filtration buffer and eluted with 10 mM maltose containing buffer. The sample were further resolved by SDS–PAGE. The experiment was repeated three times with similar result.

### GST pull down assay

The N-terminal GST tagged AMBRA1^WD40^ was co-expressed with a N-terminal TSF tag DDB1, the N-terminal GST-myc tag Cyclin D1 was co-expressed with a N-terminal TSF tag CDK4 in Expi293F cell. The complex were purified as described in the Protein expression and purification section. GST tag was remained for Cyclin D1-CDK4 complex. For GST pull down assay, 0.7uM of AMBRA1^WD40^-DDB1 complex was incubated with GST-myc-Cyclin D1-CDK4 complex in 5 to 1 molecular ratio supplemented with 50ul GST beads for 2hrs. After washing with 500ul of gel filtration buffer three times, the proteins were eluted with 100ul of gel filtration buffer supplemented with 50 mM reduced glutathione and resolved by SDS–PAGE.

### Cell culture and treatment

The human embryonic kidney HEK293 cells were cultured in Dulbecco’s modified Eagle’s medium (Gibco) supplemented with 10% fetal bovine serum and 1% penicillin/streptomycin solution at 37 °C under 5% CO_2_. The cell lines were maintained by passaging the cells, using Trypsin-EDTA solution (Gibco), after they are sub-confluent to around 75–90% in 6 cm plate (Corning). Transient transfections were performed using X-tremeGENE HP DNA Transfection Reagent according to the manufacturer’s instructions (Roche). Cells were analyzed 48 hours after transfection. For blocking autophagic flux, cells were treated with 100 nM of bafilomycin A1 for 2 hours at 37 °C.

### RNA interference

SiRNA oligoribonucleotides corresponding to the human AMBRA1 were ordered from Tsingke. AMBRA1 siRNA 1: 5’-GGCCUAUGGUACUAACAAA -3’; AMBRA1 siRNA 2: 5’-GCGGAGACAUGUCAGUAUC -3’. For siRNA transfection, a total of 4*10^5^ cells per well were plated at 6-well plate. Each well transfected with 100 pmol siRNA by using Lipofectamine RNAiMAX (ThermoFisher #13778075) next day, according to the manufacturer’s instructions. 48h after transfection, the cells were harvested for quantitative PCR reactions and re-plated into the 12-well plate for further experiment.

### Quantitative PCR with reverse transcription

Total RNA extraction was performed by using TransZol Up Plus RNA Kit (Transgenbiotech), cDNA was generated using HiScript III 1st Strand cDNA Synthesis Kit (Vazyme) according to manufacturer’s instruction. The quantitative PCR were performed using TransStart Top Green qPCR SuperMix (Transgenbiotech) with the Bio-Rad CFX Connect Real-Time PCR Detection System in a 96-well format. Bar graphs represent the relative ratios of target genes to housekeeping gene values.

### Immunoblotting and immunofluorescence

The human embryonic kidney HEK293 cells were plated into 6-well plate, then followed by SiRNA transfection using Lipofectamine RNAiMAX (ThermoFisher Scientific) next day, according to the manufacturer’s instructions. 48h after SiRNA transfection, the cells were re-plated into the 12-well plate at 8*10^4^ density per well for immunoblotting. Transient plasmids transfections were performed using X-tremeGENE HP DNA Transfection Reagent (Roche) according to the manufacturer’s instructions next day. For blocking autophagic flux, cells were treated with 100 nM of bafilomycin A1 for 2 hours at 37 °C before collection. Cells were analyzed 48 hours after transfection. Cells were lysed in lysis buffer (50mM Tris pH 7.4, 150 mM NaCl, 0.5 mM EDTA, 0.5 mM MgCl_2_, 1% NP-40) plus protease inhibitors (PMSF, aprotinin, leupeptin and pepstain). Lysates were lysed for 30min on ice, then centrifuged at 13000g for 10 min to remove insoluble debris. Solubilized proteins were quantified by BCA Protein Assay Kit (Sangon Biotech), equal amounts of protein were mixed with loading buffer and incubated at 95 °C for 5 min. whole cell lysates were separated by SDS-PAGE and transferred to 0.45 um Immun-Blot PVDF membranes (BIO-RAD). Membranes were then blocked in 5% milk/TBST for 1 hour at room temperature and incubated with primary antibodies at 4 °C overnight. For the detection of proteins, using the appropriate secondary antibodies conjugated to horseradish peroxidase (anti-mouse and anti-rabbit, Cwbio) at 1:2000 dilution in TBST for 1 hour at room temperature and visualizing with BeyoECL Plus (Beyotime). The images were acquired by Amersham Imager 680. For immunofluorescence, cells were re-plated into the 12-well plate on coverslips at 5*104 density per well 48h after SiRNA transfection, then follow the same procedure as for immunoblotting. 48 hours after plasmids transfection, cells were washed with cold PBS and fixed in 4% Paraformaldehyde (PFA) in PBS for 20 min at room temperature, then permeabilized with PBS/0.1% (v/v) Triton X-100 for 20 min and subjected to blocking with PBS/1% (w/v) BSA to block the cell for 1 hour at room temperature. Primary antibodies were diluted in PBS/0.01% (v/v) Triton X-100 at 1:1000 dilution, applied onto the coverslips at 4 °C overnight. Alexa488-conjugated secondary antibody was diluted in PBS/1% (w/v) BSA at 1:1000 dilution and incubated with the coverslips for 1 hour at room temperature. Slides were mounted in Anti-fade Mounting Medium with DAPI (Beyotime). Imaging was performed using a Zeiss LSM 900 confocal microscope with a Plan-Apochromat 63×/1.40 oil objective.

### *In vitro* ubiquitination assay

The ubiquitination assays were performed in a 25 μl reaction volume with the following components: 100 nM UBE1 (Boston Biochem E-304), 1.5 μM UBCH5C (Boston Biochem E2-627), 0.3 μM purified AMBRA1-DDB1-Cullin4-RBX1 complex, 20 μM HA-ubiquitin (Boston Biochem U-110), 0.16 μM myc-Cyclin D1-CDK4 complex and 10 mM MgATP solution (Boston Biochem B-20) in E3 ligase reaction buffer (Boston Biochem B-71). The reaction was incubated at 37 °C for 2h and analysed by SDS-PAGE, followed by immunoblot. The experiment was repeated three times with similar result.

### Quantification and statistical analyses

For western blot quantification, a minimum of three independent experiments were included in the representing graphs, band densitometry was measured by ImageJ, LC3-II levels were normalised to loading control. For LC3 puncta quantification, a puncta was defined as a LC3 positive mainly circular cytoplasmic structure of approximately 1 um in diameter, which is thought to be an autophagic vesicle. LC3 punctum were counted in a total of 50 cells in 8 different fields for each condition. Cells being counted were randomly selected and punctum were counted by using ZEN (Blue edition) software in a blinded fashion by 3 investigators. Data were analysed using GraphPad Prism 7.0.0 software. Differences between datasets were analysed using two-tailed statistical tests, the value of p < 0.05 was considered statistically significant and the conventions used to depict the results across this article are as follows: *p ≤ 0.05; **p ≤ 0.01; ***p ≤ 0.001 and Error bars represent standard deviations.

## Acknowledgments

The authors thank the cryo-EM and advanced mass spectrometry facility (KMS) of Kobilka Institute of Innovative Drug Discovery, the Chinese University of Hong Kong (Shenzhen) for the support. This work was supported by the Start-up funding from SUSTech, the Natural Science Foundation of Guangdong Province of China (to M.-Y.S., 2022A1515010856), the National Natural Science Foundation of China (to G.S., 31950410540) and Foreign Youth Talent Program from State Administration of Foreign Experts Affairs (to G.S., QN2021032004L)

## Contributions

M.L., M.-Y.S. and G.S. designed the experiments. M.L. performed the experiments. Y.W. collected the EM data. F.T., X.W., and X.M. contributed in the early stages of the project. M.- Y.S. and G.S. processed the EM dataset. M.-Y.S. and G.S. wrote the manuscript.

---

**Table S1.**
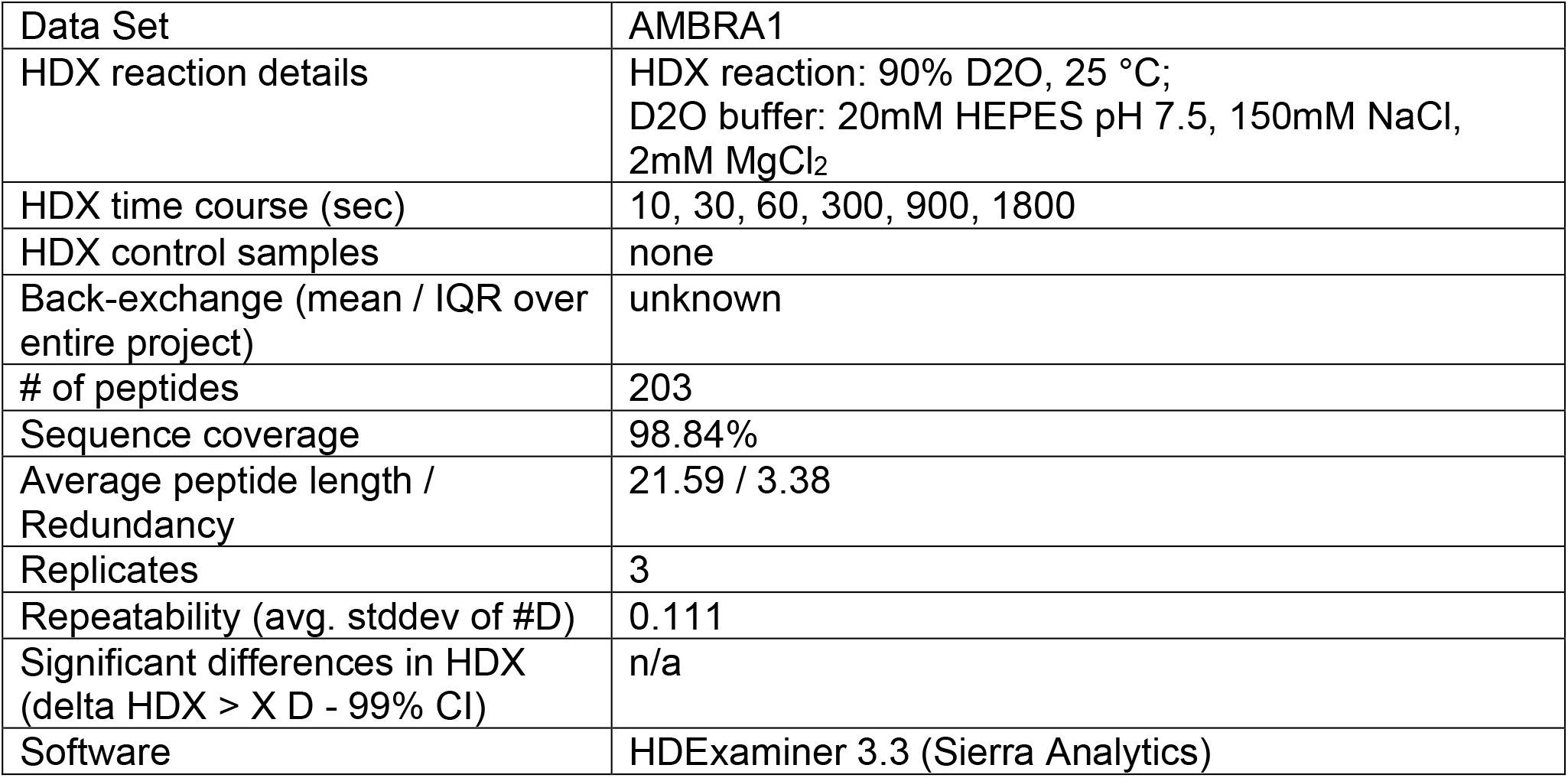
HDX.

**Table S2.**
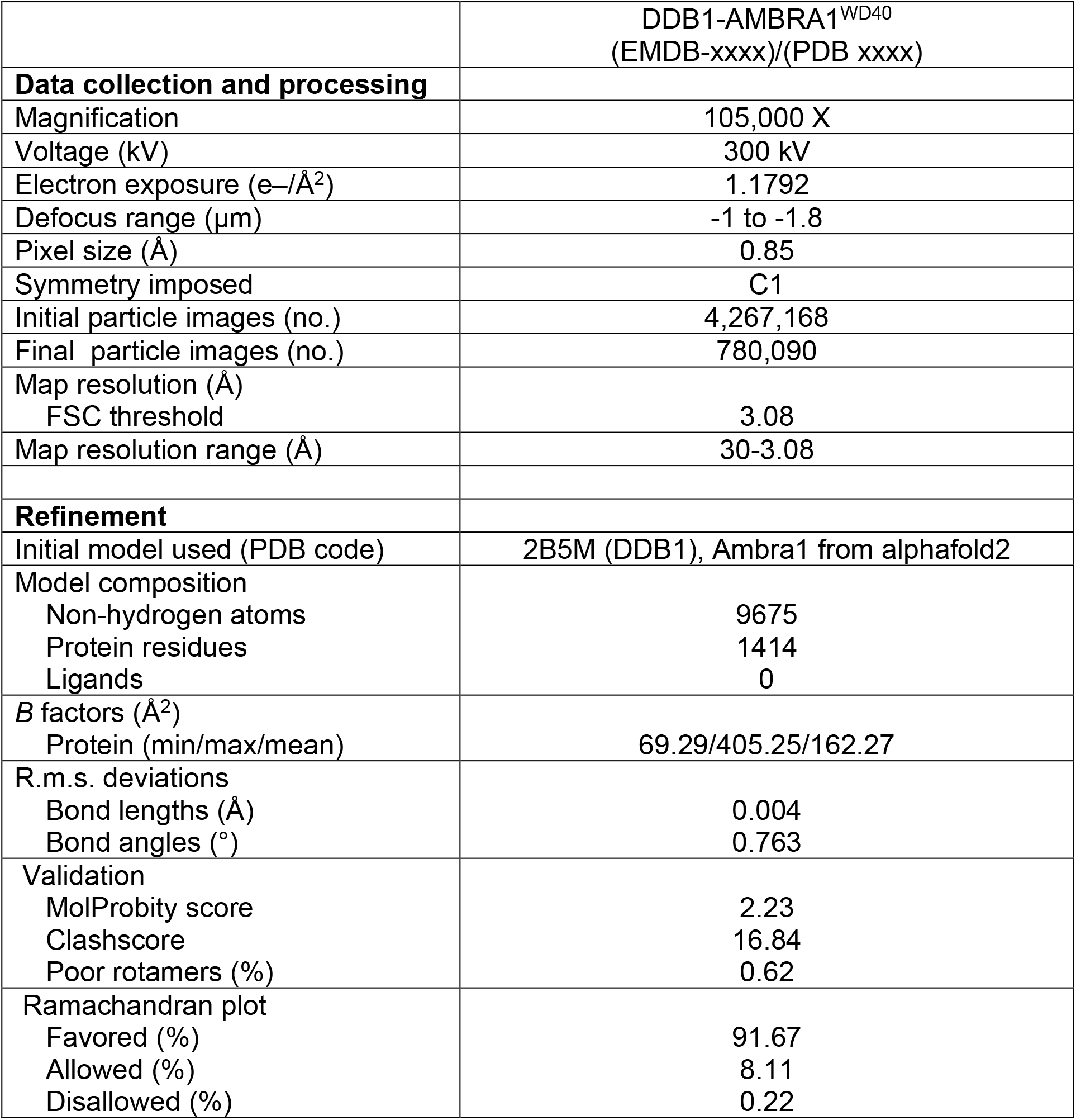
Cryo-EM data collection, refinement and validation statistics.

